# Canonical Representational Mapping for Cognitive Neuroscience

**DOI:** 10.1101/2025.09.01.673485

**Authors:** Manuel Schottdorf, Sebastian Michelmann

## Abstract

Understanding neural representations is central to cognitive neuroscience, yet isolating meaningful patterns from noisy or correlated data remains challenging. Canonical Representational Mapping (CRM) is a novel multivariate analysis method to identify neural patterns aligned with specific cognitive hypotheses. CRM maximizes correlations between multivariate datasets – similar to Canonical Correlation Analysis – while controlling for confounding sources of variance, such as shared noise or irrelevant task conditions. We validate CRM with simulations and apply it across diverse neurophysiological datasets: In one application, we use CRM to map large language model activations onto intracranial electroencephalography recordings during story listening, controlling for contextual autocorrelations. In another example, we factorize overlapping representations in functional Magnetic Resonance Imaging into distinct context and episode components. Finally, we uncover frequency-coupled representations shared between hippocampus and medial prefrontal cortex in navigating rodents. Our results introduce CRM as a powerful tool to isolate representations across modalities and species.

## Main

A central aim of cognitive neuroscience is to characterize the correlation between neural representations and cognitive and behavioral variables. For example, the finding that perceiving places and faces evokes distinct neural activity patterns^1^ can be framed as a correlation between the voxel-wise activation patterns and categorical task variables (“place” vs. “face”). In practice, neural, cognitive, and behavioral variables are often multivariate^2,3^, requiring analytical tools that capture the relationships between two multivariate series of observations; the rise of multivariate analysis methods in neuroscience spurred a wide range of methods for studying representations. These include techniques for mapping brain-behavior associations (e.g.,^4–6^), characterizing inter-regional connectivity^7–9^, and aligning neural activity across individuals (e.g.,^10–12^).

A unifying assumption across these methods is that similar representations are characterized by similar patterns of activity. Representational similarity analysis (RSA) formalizes this idea by studying the pairwise similarity structure of multivariate patterns – summarized in a representational dissimilarity matrix (RDM)^13,14^. By contrast, alignment techniques take the complementary approach: transforming multivariate patterns such that the resulting components are maximally aligned – by components we mean the synthetic variate that is derived as a linear combination of features. For instance, multi-subject fMRI data can be succinctly described by a reduced number of orthogonal components in a shared response model^15^. One widely used method for multivariate alignment is Canonical Correlation Analysis (CCA)^16,17^. CCA identifies linear transformations for each of two multivariate datasets such that the resulting canonical variates (components) are maximally correlated.

Importantly, not all correlations between multivariate measurements reflect meaningful representational overlap. Alignment methods that maximize correlations can inadvertently capture trivial or artifactual activity such as electrical noise or scanner drift. Moreover, neural representations often co-occur with other representational patterns that are not of primary interest. One recent example of this problem is that the neural representations of individual words in a narrative are inherently entangled with the broader contextual structure of the story. When aligning word-embeddings derived from Large Language Models (LLMs) with neural activation, the resulting shared representations may consequently capture those unwanted aspects^18,19^. Furthermore, neural representations that are computationally important may only explain a small amount of variance in neural measurements^20^, requiring transformations of the data that isolate representations of interest from overlapping representational patterns. In one recent functional Magnetic Resonance Imaging (fMRI) study^21^, overlapping representations were observed during repeated viewing of naturalistic movies. In those movies two protagonists visited two contexts from two different schemata (cafés and supermarkets). Patterns of neural activity in medial PFC showed stronger evidence for schematic than for contextual or episodic representations; nonetheless, contextual representations in mPFC may exist, and researchers may wish to isolate and study them. These challenges motivate the need for a method that can factorize multivariate patterns into separate representations, effectively maximizing a similarity structure that is derived from a cognitive hypothesis (Figure 1), while controlling for similarity from sources of variance not of interest.

**Figure 1:**
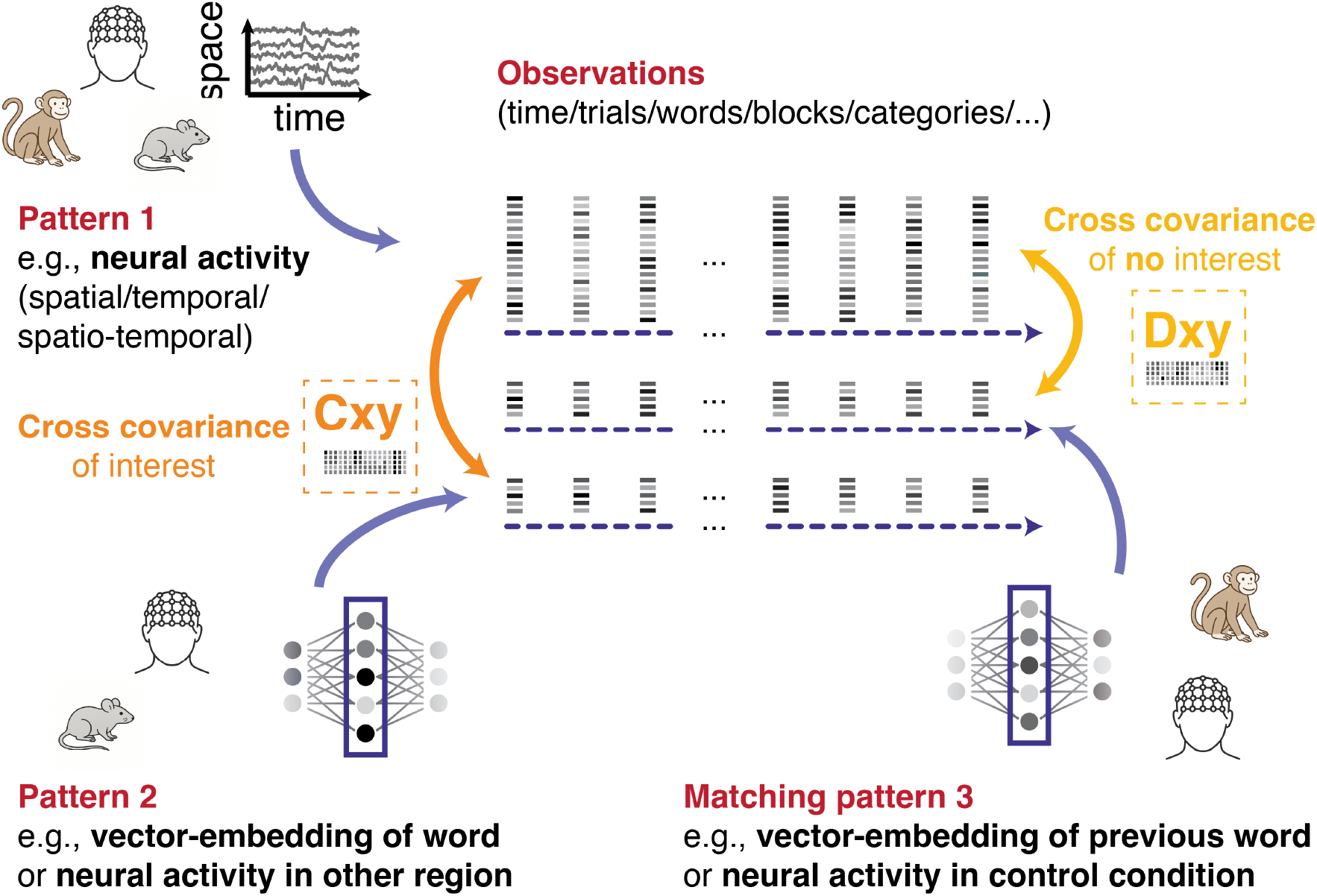
Motivation for Canonical Representational Mapping. Shown is a conceptual overview of CRM: When researchers collect two observations with repeated measures (e.g., any pair of continuous multi-channel recordings from different brain regions in human or animal recordings, different subjects, vector embeddings in neural network models, or multivariate behavioral data, etc.) they can ask if the same information is represented in both time series. Three example multivariate series of observations are illustrated (red labels). If the relationship between pattern 1 and pattern 2 is of interest, the pairwise linear relationships can be captured in the cross-covariance matrix between the multivariate time series (*C*_*xy*_, orange). Canonical correlation analysis (CCA) can transform them, so that resulting components are maximally correlated. However, shared information can stem from confounding variables and common noise sources. A third series of observations (pattern 3) may share the confounding signal of no interest and can be used to estimate the pairwise linear relationships of confounding signal between the variates (*D*_*xy*_, yellow). Canonical Representational Mapping leverages this control-covariance to separate neural representations from noise and disentangle overlapping representational patterns. I.e., akin to CCA, CRM maximizes the correlation between two multivariate time series; however, it constrains the covariance with a matched control time series (*D*_*xy*_) to zero. Examples of control time series are lag-shifted data (e.g., a time-series of word-embeddings of the previous word, which shares the same context but not the same content), data filtered in a noise band (e.g., data that has been band-pass filtered around the line-noise), or pseudo-randomized data, where the correspondence of interest is shuffled but the correspondence of confounds is kept (e.g., trials are shuffled within a block to separate item representations from block-specific effects).

We propose a new method that solves this problem as a constrained alignment problem: Canonical Representational Mapping (CRM) maximizes the correlation between two multivariate series of observations (*C*_*xy*_), while controlling for correlations that are not of interest. This is accomplished by constraining the covariance with a matched control time series (*D*_*xy*_) to zero. We first outline the theory behind this method and then test its effectiveness through simulations and multiple applications to neurophysiological data. In one analysis, we map activations of a large language model (i.e., word-embeddings) onto intracranial Electroencephalography (iEEG) recordings (akin to Goldstein et al.^18^). Using CRM we control for contextual information by fixing the correlation with a lag-shifted time-series of embeddings (i.e., correlation with the previous word’s embedding) to zero. This analysis confirms the presence of word-representations prior to word onset while ruling out alternative explanations based on stimulus-stimulus or contextual correlations. In another analysis, we factorize the fMRI signal from Reagh et al.^21^ into components that capture separate aspects of overlapping situations (e.g., context and episode patterns). In a last example, we demonstrate the usefulness of CRM for detecting spectral features in the LFP recordings in freely behaving rodents solving a complex maze. We identify shared representations between hippocampus and medial prefrontal cortex^22^ and demonstrate that maximizing correlations via

CCA latches on to shared noise in the recording: The resulting CCA components predominantly reflect the correlation of PSD at 60 Hz; with CRM, we control the covariance in the band-pass-filtered data and detect coupling in the theta and beta frequency bands in a data-driven way.

Our results show that CRM can be a powerful tool for cognitive neuroscience to isolate meaningful neural representations across modalities and species. We end by discussing the limitations of CRM, and propose potential future extensions.

### Methodological validation and simulation

While full mathematical details are described in the Methods section, we wish to develop an intuition for how CRM works. Readers primarily interested in neurophysiological applications may skip ahead to the following sections. In brief, CRM maximizes the correlation corr(*w*_*x*_*X, w*_*y*_*Y*) over the weight vectors *w*_*x*_ ∈ ℝ^*p*^ and *w*_*y*_ ∈ ℝ^*q*^, given the data *X* ∈ ℝ^*p×n*^ and *Y* ∈ ℝ^*q×n*^ where *p, q* is the number of variates, and *n* is the number of observations. These data typically take the form of variates over observations, for example BOLD or EEG signals measured over time (or trials) across many voxels or electrodes. CRM maximizes this correlation under the constraint that applying the same weights to datasets *S* ∈ ℝ^*p×n*^ and *T* ∈ ℝ^*q×n*^ yields corr(*w*_*x*_*S, w*_*y*_*T*) *≈* 0. For example, *S* and *T* could be EEG signals during inter-trial intervals, while *X* and *Y* are taken during trials (Fig. 1). Alternatively, the same data can be reorganized so that *S* and *T* lose only the key association of interest (e.g., trial correspondence) while retaining associations not of interest (e.g., block correspondence). This formulation can be restated as a quadratic optimization problem on the covariance matrices *C*_*xy*_ = cov(*X, Y*) and *D*_*xy*_ = cov(*S, T*), aiming to maximize *w*_*x*_*C*_*xy*_*w*_*y*_ while enforcing *w*_*x*_*D*_*xy*_*w*_*y*_ = 0 with a Lagrange multiplier. This constrained quadratic problem yields many solutions for *w*_*x*_ and *w*_*y*_, computable in polynomial time via eigendecomposition (see Methods).

To develop an intuition for CRM solutions, specifically in comparison to CCA, we performed a set of analyses on simulated data. In these simulations, we controlled the correlation level by generating *X* ∈ ℝ^*p×n*^ from normally distributed random numbers with unit variance, and deriving *Y* = *M × X* + *α η* where *M* ∈ ℝ^*q×p*^*∼* 𝒩 (*µ* = 0, *σ* = 1) is a fixed matrix with random entries drawn from a normal distribution with unit variance. The contribution of normally distributed noise *η* ∈ ℝ^*q×n*^ was set with a noise parameter *α*. This procedure yielded the correlated datasets *X* and *Y*. We additionally simulated two further random datasets *S* ∈ ℝ^*q×n*^ *∼* 𝒩 (*µ* = 0, *σ* = 1) and *T* = (*βN* + (1 − *β*)*M*) *× S* where *N* ∈ ℝ^*q×p*^ *∼* 𝒩 (*µ* = 0, *σ* = 1) is a random matrix, and *M* is the same as above. The partial alignment of *X, Y* and *S, T* via the shared matrix *M* makes the CRM problem challenging. The parameter *β* tunes this alignment, and was set ad-hoc to *β* = 1*/*4.

In the absence of noise, *α* = 0, the optimal CCA weights satisfy *w*_*x*_*Mw*_*y*_ = 1 and *w*_*x*_*C*_*xy*_*w*_*y*_ = 1. Computing *w*_*x*_*D*_*xy*_*w*_*y*_ in this setting yields some finite value, as CCA does not account for the constraint. CRM, in contrast, enforces *w*_*x*_*D*_*xy*_*w*_*y*_ *≈* 0 across all noise levels (Fig. 2a,b). Of note, we display the results with the strongest correlation *w*_*x*_*C*_*xy*_*w*_*y*_ because multiple solutions exist for both methods; for CCA, their number is determined by the number of existing eigenpairs. We next sought to illustrate the various solutions of CRM. In CRM, valid solutions are roots of the eigenvalues of a matrix as function of a tuning parameter *λ*_3_ (see Methods). Plotting the eigenvalues of our simulated problem for one example noise level as function of this tuning parameter reveals numerous zero crossings (red dots in Fig. 2c) where the constraint is satisfied. Each zero crossing corresponds to a CRM solution, generally with a different correlation value *w*_*x*_*C*_*xy*_*w*_*y*_. To illustrate this, we color-coded the curves according to their correlation values. Note that the different curves all flatten out towards larger values of *λ*_3_; as shown in the Methods section, no solutions exist for large *λ*_3_. Consequently an exhaustive search can find all CRM solutions, and the best solution can be identified based on its largest eigenvector. Example implementations of both methods are provided as Matlab and Python code in our repository (see: Code availability).

**Figure 2:**
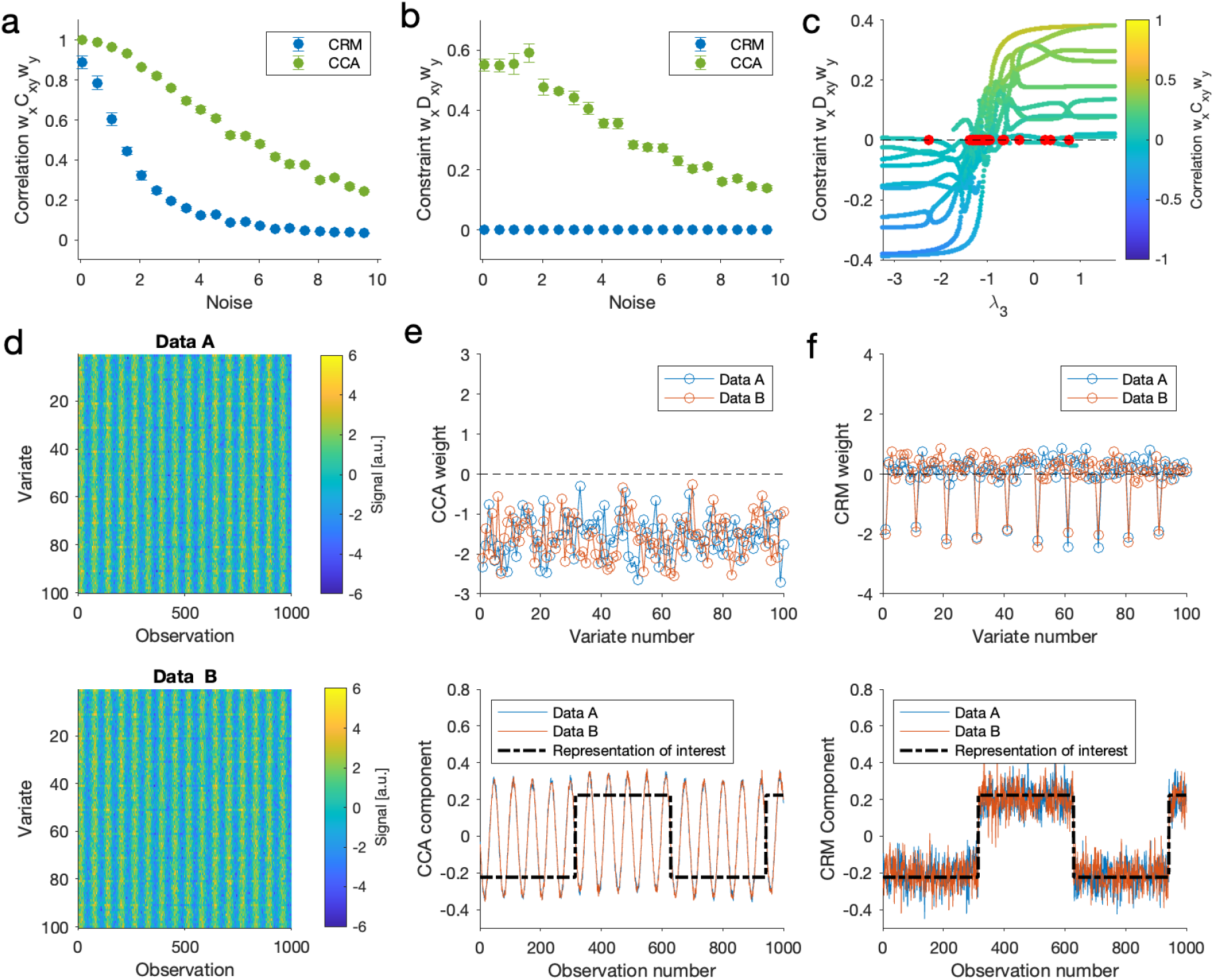
Proof of principle of CRM on simulated data. **a** Best correlations produced by CRM (blue) and CCA (green) for two random matrices *X* and *Y*, where *X* is populated by normally distributed random numbers, and *Y* = *M × X* + *α η* is a rotated version of *X*, corrupted by noise of magnitude *α*. Values are means over *N* = 10 repeats, error bars indicate S.E.M. **b** Projection *w*_*x*_*D*_*xy*_*w*_*y*_ of CCA and CRM weight vectors *w*_*x*_ and *w*_*y*_ onto the constraining matrix *D*_*xy*_ for various noise levels *α* for the data in (a). The vectors produced by CCA produce finite values, whereas CRM enforces *w*_*x*_*D*_*xy*_*w*_*y*_ *≈* 0. **c** *w*_*x*_*D*_*xy*_*w*_*y*_ as a function of *λ*_3_ for an example dataset. Curves are color-coded by the corresponding correlation *w*_*x*_*C*_*xy*_*w*_*y*_; red dots mark points where the constraint is met. Note how curves flatten out towards large *λ*_3_; all solutions lie in the center of the plot. **d** Simulated dataset containing a representation of interest (square wave) embedded in a subset of variates, masked by a shared sinusoid. **e** Top: CCA weights (blue/orange) for the data in (d, top/bottom), showing roughly uniform weighting across channels. Bottom: CCA-components compared to the signal of interest (black dashed). The CCA components (orange/blue) are dominated by the sinusoidal noise. **f** Top: Same as e, but with weights computed with CRM. Note how most weights scatter around 0, with large weights restricted to variates containing the target representation. Bottom: components computed with CRM; components closely match the target signal (black dashed), indicating successful isolation of the representation of interest.

We next simulated a neuroscience-motivated use-case (Fig. 2d-f). We generated multiple observations of a set of variates containing a representation of interest (a square function) that was obscured by other signals, here modeled as a high-amplitude sinusoid. The square signal is linearly added to every 10th variate; thus the representation of interest was present only in variates 0, 10, 20, and so on. In addition to this shared representation masked by the shared sinusoidal noise signal, two simulated datasets *X* and *Y* (Fig. 2d) also contained uncorrelated noise unique to each variate and observation. From these two unfiltered datasets, we computed the covariance *C*_*xy*_. To extract the slow representation of interest, we band-pass filtered the data in the time domain, isolating the noise frequency. From this noise-band-filtered data we computed the constraining covariance *D*_*xy*_, which was then used to constrain the alignment of the original unfiltered data (see Methods).

As expected, CCA alignment produced canonical coefficients (Fig. 2e top) dominated by the sinusoidal signal, which was unrelated to the representation of interest (Fig. 2e bottom). In contrast, CRM assigned large weights only to variates containing the representation of interest (Fig. 2f top). The resulting CRM components closely matched the target signal (see Fig. 2f bottom), indicating successful isolation of the representation of interest.

### Context-free alignment of neural representations and vector embeddings

Several recent studies have shown that the vector representation generated by large language models (LLMs) — i.e., word embeddings — resemble the information represented in human neural activity^4,18,23^. For example, Goldstein et al.^23^ fit regression models mapping word-embeddings onto high-frequency activity in human iEEG during naturalistic story listening. In their study, cross-validated encoding models predicted neural activity from word embeddings, indicating that LLMs and humans represent word information in similar ways. A key finding was the presence of word representations in neural activity prior to the onset of the word, which the authors interpreted as the representation of upcoming words via neural mechanisms of prediction in natural language. A recent study, however, criticizes these findings and suggests that stimulus dependencies rather than prediction explain the fit of encoding-models prior to word-onset^19^; that is, correlations between the current word representation and the previous word representation may give rise to spurious pre-stimulus alignment. This is an exemplary case of why controlling confounding correlations is essential when aligning representations between systems.

Indeed, correlations between multivariate patterns in word embeddings and human neural activity can arise from both context and content representations, since both are captured by the embeddings and the neural data. To control for this confound, we reasoned that – if the context-representation of the current word is also captured by the previous word – we could remove its influence by restricting the alignment with the previous word embedding. Accordingly, we computed the channel-by-embedding cross-covariance matrix *C*_*xy*_ across words in the story to maximize correlations between neural activity and the top 50 principal components of the embeddings (see Methods). For CRM, we additionally computed *D*_*xy*_ — the cross-covariance between neural activity and the embedding of the previous word (Fig. 3a) — and enforced *w*_*x*_*D*_*xy*_*w*_*y*_ = 0 during alignment. We repeated this analysis at lags from -4 s to +4 s around word onset. For comparison, CCA was computed by setting *D*_*xy*_ to zero. The correlation of the first CCA and CRM components on held-out data is shown in Fig. 3b. Both methods achieved high correlations (*>*.3) on the test set, increasing prior to word onset. Crucially, only correlations between the CRM components decayed fully to zero within the 4 seconds before word onset. Specifically, 1.5 seconds before word onset the maximization of correlations in the training set no longer generalized to the test set when the correlation with the previous-word embedding was constrained to zero. This suggests that pre-stimulus correlations of the CRM component can be interpreted as prediction of the upcoming word (content prediction), whereas the CCA component may capture correlations with context-representations that are shared across adjacent words. Component loadings on channels (Fig. 3c) suggest a broadly similar distribution of the neural representation for CCA (Fig. 3c), top) and CRM (Fig. 3c), middle), with higher loadings for CCA in early auditory processing regions and higher loadings for CRM in higher-order auditory processing regions (e.g., posterior STS and right IFG, Fig. 3c), bottom). This is consistent with the interpretation that stimulus-driven correlations in early auditory areas can influence the CCA results.

**Figure 3:**
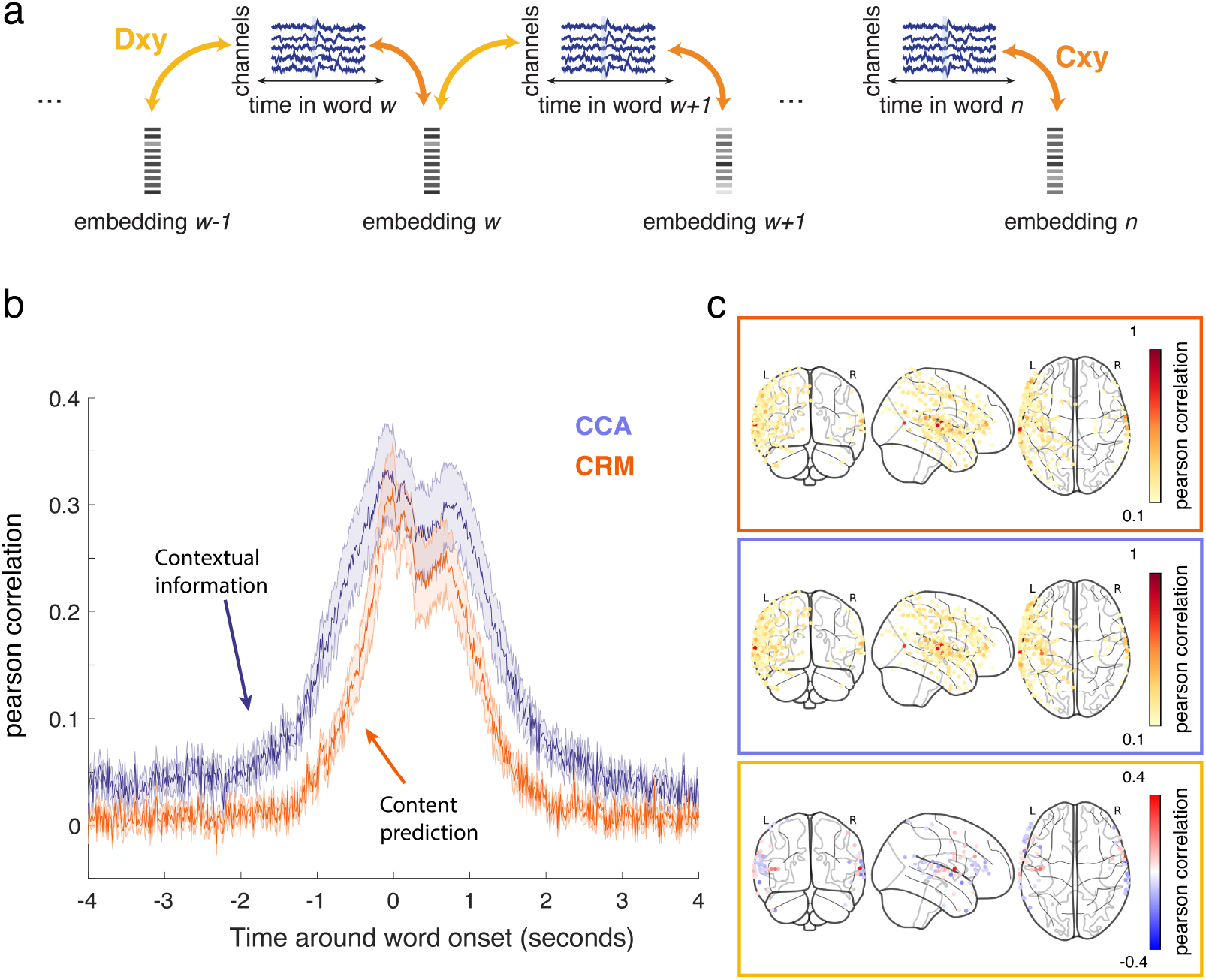
Alignment between vector embeddings and intracranial EEG recordings in humans. **a** Vector embeddings from Whisper^24^ are mapped onto high-frequency activity in iEEG during naturalistic story listening^25^. Both CRM and CCA derive component pairs that maximize the cross-covariance between vector embeddings and neural activity *w*_*x*_*C*_*xy*_*w*_*y*_. Only CRM additionally constrains the cross-covariance between neural activity and the vector embedding of the previous words to zero (*w*_*x*_*D*_*xy*_*w*_*y*_ = 0). **b** Mean Pearson correlation *±* SEM (across 9 patients) in held-out data for CCA (purple) and CRM (orange), averaged across 10 folds. Correlations were computed at lags relative to word onset. CCA yields positive correlations up to 4 s before onset, potentially reflecting prediction of upcoming words and/or shared contextual information between embeddings within the same story. CRM yields positive correlations from *∼* 1 s before up to *∼* 2 s after onset, indicating that word-specific representations are present in neural activity during this window. Only the CRM-derived correlations can be interpreted as predictions of word-level representations. **c** Component loadings on iEEG channels for CCA (top), CRM (middle), and their difference (bottom). Warm colors in the difference map indicate stronger contribution to CCA; blue indicates stronger contribution to CRM. Only channels where the absolute difference exceeded 0.05 are plotted.

Of note, the linearity of CRM is advantageous in analyses of these types, because a sufficiently powerful non-linear method can learn to predict the next word based on the statistics of natural languages. In other words, the increase of correlations before word onset in Fig. 3b is not due to CRM learning to predict the word from neural activity, but rather to the fact that upcoming content can be linearly decoded from neural activity. A more complex scheme would make this analysis harder to interpret (see also Discussion).

### Factorizing overlapping representations into their constituent parts

A plethora of studies has investigated how information is represented in different brain regions^13^. One recent fMRI study systematically manipulated stimulus material to probe neural representations of nested properties of naturalistic experience^21^. In 24 different videos, two characters (Tommy and Lisa) visited four locations belonging to two schemata (café and grocery store). Among other findings, the authors observed that medial prefrontal cortex (mPFC) held schematic representations: pairwise correlations between individual trials were most consistent with a model in which videos from the same schema elicited similar neural response patterns (Figure 4a). Other models – based on shared context (e.g., patterns of Tommy in cafe 1 correlating with Lisa in cafe 1), or shared episode (e.g., pattern correlation between three videos of Tommy visiting café 1)– fit the data to a lesser extent. This motivates the possibility that overlapping representations in mPFC can be factorized into constituent parts, potentially isolating contextual and episodic representations that are masked by dominant schematic representations.

**Figure 4:**
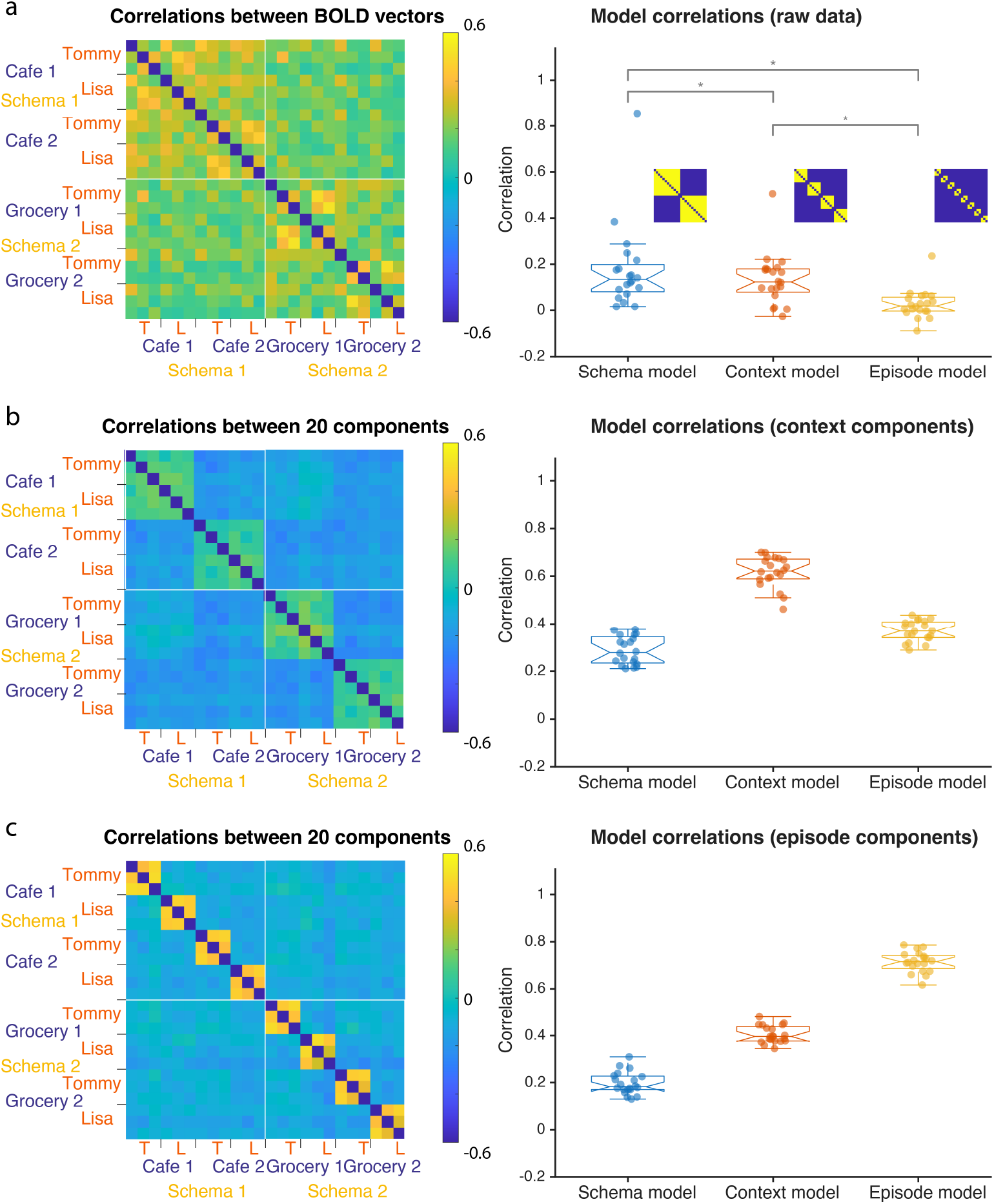
Representational Similarity Analysis (RSA) with CRM. CRM can factorize overlapping neural representations into their constituent parts. **a** RSM of fMRI BOLD patterns in medial prefrontal cortex (mPFC; data from Reagh and Ranganath, 2023^21^). Participants viewed three videos of four contexts visited by two individuals. Contexts belonged to two schemata (café and grocery store). Multivariate BOLD responses correlate between the trials; correlation of the RSM with a schematic model matrix was significantly higher than with context or episode models suggesting that mPFC predominantly holds schematic representations. **b** Correlation across 20 CRM “context” components: CRM was fitted to maximize the cross-covariance between patterns of the same context (here: *M*_*xy*_) while keeping the cross-covariance between patterns of different context in the same schema (*L*_*xy*_) at zero. Correlations between the RSM and a context model-matrix are strongest. **c** Correlation across 20 CRM “episode” components: CRM was fitted to maximize the cross-covariance between patterns of the same situation (same person in the same context; *N*_*xy*_) while keeping the cross-covariance between patterns of different situations (different person in the same context; *M*_*xy*_) at 0. Correlations between the RSM and an episode model-matrix are strongest.

We first reproduced the original finding by computing a Representational Similarity Matrix (RSM) as the pairwise correlation between beta weights from all videos (Figure 4a, left). Correlating the RSM with a schematic model matrix (where in-schema correlations are 1 and other correlations are 0) yielded the highest average correlation across 20 subjects (*r* = 0.177, SD = 0.185). This correlation was significantly higher than the correlation with the contextual model matrix (*r* = 0.135, SD = 0.1137, *p <* 0.04) and the correlation with the episodic model matrix (*r* = 0.027, SD = 0.064, *p <* 0.001). The correlation with the contextual model matrix was also significantly higher than the correlation with the episodic model matrix (*p <* 0.001, Figure 4a, right).

We then transformed voxel-wise covariance into component representations aligned with contextual and episodic models. To do this we computed the pairwise linear relationships between voxels across different trial-correspondences: To capture schema-level structure, we computed the covariance matrix *L*_*xy*_ from trial pairs that shared a same schema but differed in context (e.g., grocery store 1 and grocery store 2 pairs). We further computed the covariance matrix *M*_*xy*_ from trial-pairings within the same context (e.g., videos of Tommy and Lisa visiting cafe 1). Finally we computed the covariance matrix *N*_*xy*_ from trial-pairings within the same situation (e.g., the three videos of Tommy visiting grocery store 1). Using the fast solver (Algorithm 2) we derived 20 components that maximized correlations in the same context *M*_*xy*_ while the correlations between trials that formed part of the same schema but not the same context (*L*_*xy*_) to zero. We then re-computed the RSM across these 20 components, i.e., we compared the pattern-correlation across components for each pair of trials (Figure 4b, left) and arranged them in the same matrix-format as for the RSM of the raw data. We then correlated this RSM with the schematic, contextual, and episodic model matrices. Confirming the successful transformation of the feature-data, the correlation with the context model was highest for the context-components (*r* = 0.618, SD = 0.0645); correlations with the schema-model (*r* = 0.288, SD = 0.059) and the episode-model (*r* = 0.37, SD = 0.044) were substantially lower (Figure 4b, right).

Finally, we derived 20 episodic components by maximizing the correlations in *N*_*xy*_ while constraining correlations in *M*_*xy*_ to zero. We re-computed the RSM across these 20 components (Figure 4c, left) and correlated it with the schematic, contextual, and episodic model matrix. For these episode components, the correlation with the episode model was now highest (*r* = 0.714, SD = 0.043), followed by correlations with the context-model (*r* = 0.404, SD = 0.036) and the schema-model (*r* = 0.197, SD = 0.047, Figure 4c, right). Together, these results demonstrate that overlapping representations can be factorized into their constituent parts using CRM.

### Unsupervised discovery of coupled spectral features in LFP recordings

In electrophysiological recordings, shared noise is common, often arising from 50 Hz or 60 Hz line interference. This can make the unsupervised search for relationships across wide spectral bands challenging. To demonstrate this, we analyzed previously published data from medial prefrontal cortex (mPFC) and Hippocampus (HPC) of awake, behaving rats engaged in a complex spatial navigation task^26^. Spectrograms from the electrode time series revealed strong correlated noise within a narrow frequency band (Fig. 5a,b). This dataset was recorded in the United States, where the line frequency is centered at 60 Hz. Applying CCA to survey the spectra for correlated fluctuations across frequencies, produced large weights around the 60 Hz band. To maximize correlations while controlling for this noise, we filtered the data with a band-pass filter around 60 Hz to obtain matrices *S* and *T*, and applied CRM to keep the correlation in this noise-band at zero. This procedure removed the strong line-noise bias, and revealed significant correlations in the theta band with minor beta contributions (Fig. 5c). The CRM components over time produce a time series dominated by fluctuations on the time scale of seconds; the same time scale as the animal navigating in the maze. This suggests that the slowly evolving cross-area correlations in the theta and beta bands are related to behavior (Fig. 5d).

**Figure 5:**
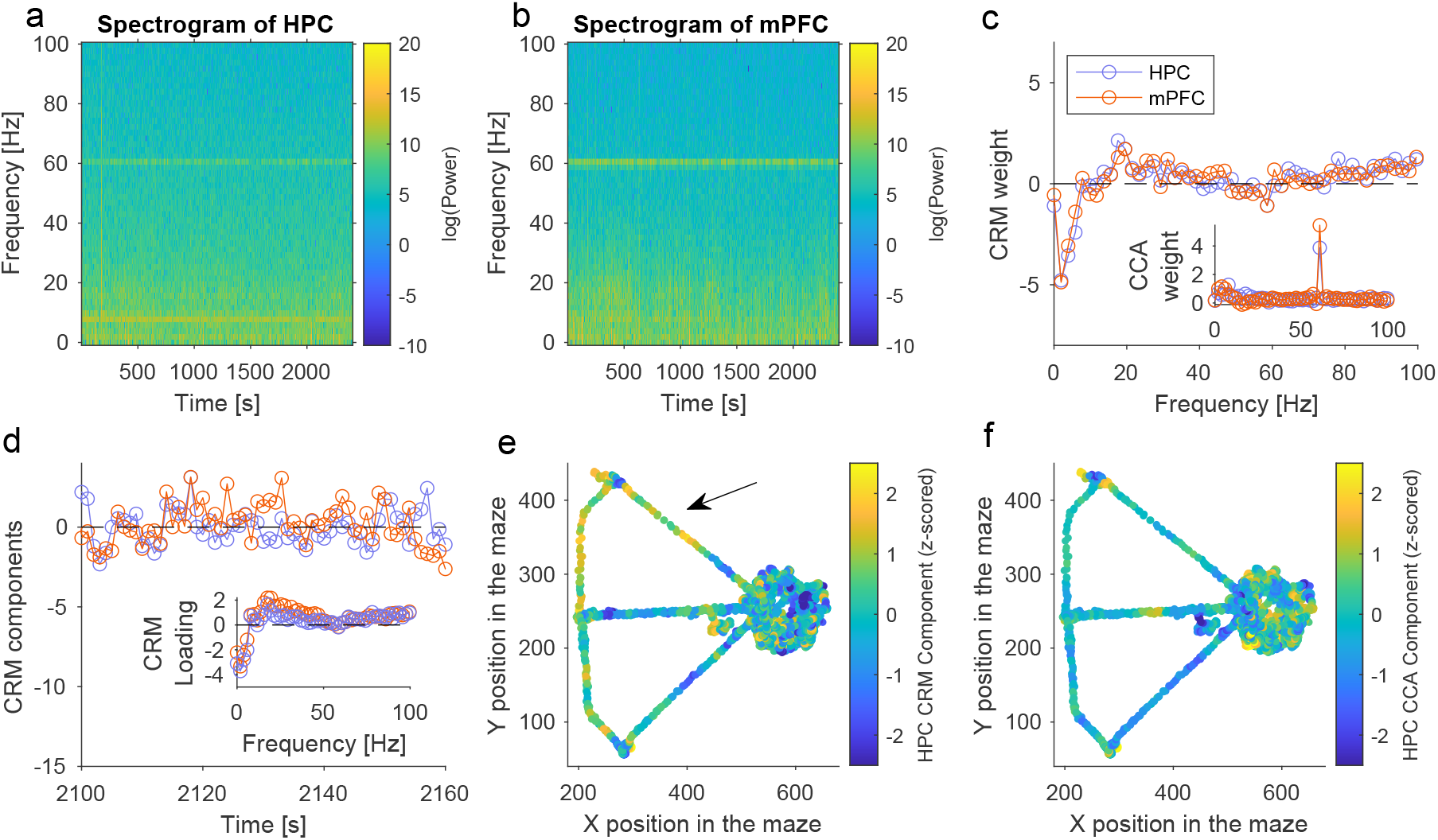
Unsupervised discovery of spectral features in LFP recordings in the presence of noise in rodents navigating a maze. **a,b** Spectrograms of LFPs recorded in the mPFC and HPC of awake and behaving rats solving a spatial navigation task^26^. **c** Weights computed with CRM for the data shown in a/b in (blue/orange). Insets show the same weights computed with CCA. Note that the CCA weights peak at a shared noise frequency around 60 Hz. **d** CRM-components from the data in a (blue) and b (orange) as a function of time. The inset shows the spectral loadings from a/b, indicating the largest amplitude-amplitude coupling in the theta band with minor beta contributions. **e** HPC CRM component from d as a function of position in the maze. Warmer colors appear away from the home box of the animal (black arrow); the home box is located at X position = 600 cm and Y position = 250 cm. **f** Same as e, but components computed with CCA. No relationship with the maze-position is apparent, consistent with CCA components being dominated by 60 Hz noise.

Plotting the CRM components against the animal’s physical position in the maze revealed an interpretable pattern: the CRM components (approximately a weighted difference between log-theta and log-beta power) were small in the home box and increased away from the home box and towards the choice point and in the reward sections of the maze. This pattern was similar in mPFC and HPC. Warmer colors away from the home position in Fig. 5e illustrate this effect. We quantified this relationship with the Pearson correlation coefficient between distance from the home box and the CRM component (*r* = 0.1); compared to random permutations, this was highly significant (*p* = 0.002). By contrast, the CCA component had no relationship with distance to the home box (*r* = −0.04; *p* = 0.2); plotting the CCA components as a function of the physical position of the rat in the maze did not reveal any relationship (Fig. 5f), consistent with the fact that CCA-based coupling was dominated by 60 Hz line noise. The examples shown in Fig. 5 display the HPC CRM component. The mPFC CRM component (not shown) was statistically indistinguishable. Although in practice, electrophysiologists often target frequency bands based on a priori hypotheses^26^, this result demonstrates how coordinated amplitude-amplitude fluctuations across brain areas can be identified in an unsupervised search.

## Discussion

Patterns of neural activity that align with cognitive constructs and behaviors serve as key explanatory variables at the physiological level in cognitive neuroscience. Indeed, task related neural activity can be productively described within a framework in which operations or computations transform between representations^27–29^. Here we introduced a new multivariate method to isolate representations that align with specific cognitive hypotheses. Canonical Representational Mapping (CRM) maximizes the correlation between two multivariate series of observations while constraining the correlation with matched control data to zero. Conceptually, CRM can be understood as a multivariate extension of a partial correlation: While CCA implements a multivariate extension of correlation that does not control for confounding variables^30^, CRM controls correlations specified in the covariance *D*_*xy*_.

We demonstrated that this method can solve various challenges in cognitive neuroscience data analysis. In one application, we separated neural representations predicting upcoming words in a story from those reflecting shared context^18,19,23^. Several recent papers have aligned LLM-derived representations with human iEEG recordings, identifying neural signatures of next-word prediction. However, disentangling prediction from shared context has remained challenging. One recent paper suggests that rather than reflecting signatures of next-word prediction, shared representations between LMMs and iEEG prior to word onset can be succinctly explained by stimulus-dependencies^19^. By controlling for shared context representations with CRM, we showed that this is not the case and that neural patterns uniquely characterizing the upcoming word are indeed represented before word onset.

Another challenge that CRM addresses is isolating neural representations that correspond to a specific cognitive construct. In fMRI bold signal from human mPFC^21^ we demonstrated that – despite schematic structure dominating representational similarity – contextual and episodic representations can be found via linear transformation of the multivariate response patterns. This type of transformaion via CRM allows for the targeted investigation of pre-specified neural representations: CRM can factorize overlapping representations into their constituent parts, isolating neural activity patterns that uniquely reflect one construct of interest.

Finally, similar information may also be represented in different brain regions. However, transformations aligning series of observations between regions are notoriously impacted by shared noise; multivariate alignment may easily pick up on this noise. Estimating this shared noise – for instance by band-pass filtering the data around 50 or 60 Hz – makes it possible to constrain the alignment with CRM such that correlations between noise are not part of the aligned signal.

It is noteworthy that we only discussed CRM with the constraint of *w*_*x*_*D*_*xy*_*w*_*y*_ = 0, effectively controlling for correlations that are not of interest. However, the constraint can easily be defined more generally based on known correlations. I.e., controlling correlations at a predefined target value *r* yields the constraint *w*_*x*_*D*_*xy*_*w*_*y*_ = *r* for any *r* ≠ 0.

As an extension of CCA, CRM can leverage techniques developed to improve CCA performance. For instance, “sparse CCA” imposes constraints on the weights of CCA directly, regularizing *w*_*x*_ or *w*_*y*_ with an 𝓁^1^ norm. Several examples of this and other linear constraints can be found in the literature^31–34^. Covariance matrix regularization can also be combined with CRM. When dealing with large numbers of variables, the within-set covariance matrices (*C*_*xx*_ and *C*_*yy*_) are also large and can be of the same order of magnitude as the number of observations. If the number is smaller than the number of features, the covariance matrices cannot be estimated reliably from the sample. In these cases, the covariance matrices *C*_*xx*_ and *C*_*yy*_ are usually replaced by identity matrices, and sparse CCA becomes equivalent to sparse Partial least squares^35^. In this regime regularization of the covariance matrices can also prevent overfitting^36^. In regularized CCA, identity matrices are added to covariance matrices to improve robustness in low-sample regimes^34^. Similarly, it may be desirable to add regularization to the covariance matrices *C*_*xx*_ and *C*_*yy*_ used in CRM to prevent overfitting.

Although CRM solves a quadratic constraint, the resulting transformations of the data are linear. Linear transformation offer advantages in data efficiency and interpretability (see Ivanova et al.^37^ for an in-detail discussion of this topic). It is also noteworthy that a similar conceptual idea of bringing exemplars closer to positive exemplars of the same class than to a negative example of a different class by at least a certain margin is captured in the learning objective defined by triplet loss^38^. In other words, deep neural network models with a triplet loss learning objective can find non-linear mappings that capture a related idea to the concept of CRM. To extend CRM to non-linear alignment problems, a nonlinear extension can be formulated by applying a “Kernel Trick” on the features^39^, projecting them into a high-dimensional space before applying CRM. Similarity, a deep version akin to deep CCA (dCCA) and related methods^9,40^ could nonlinearly extend CRM.

Our findings and simulations demonstrate CRM to be a flexible, and extensible method with broad potential to advance cognitive neuroscience by bridging between neural recordings and cognition. It is a novel tool designed specifically to study neural representations within and across information processing systems. While introduced with applications in cognitive neuroscience in mind, we note that CRM can have broader applications, akin to CCA, which has found wide use across economy, ecology, and genomics^33,41–43^ – CRM may prove valuable beyond neuroscience.

## Methods

### The CRM optimization problem and numerical solution

CRM aims to maximize the correlation between two multivariate datasets within a constrained subspace. Formally, the optimization problem is defined as follows:

Given two sets of observations, *X* ∈ ℝ^*p×n*^ and *Y* ∈ ℝ^*q×n*^, we seek weight vectors *w*_*x*_ ∈ ℝ^*p*^ and *w*_*y*_ ∈ ℝ^*q*^ that maximize the correlation 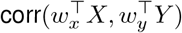, subject to the constraint that 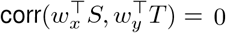 between the sets of observations, *S* ∈ ℝ^*p×n*^ and *T* ∈ ℝ^*q×n*^.

The optimization objective can be written as:

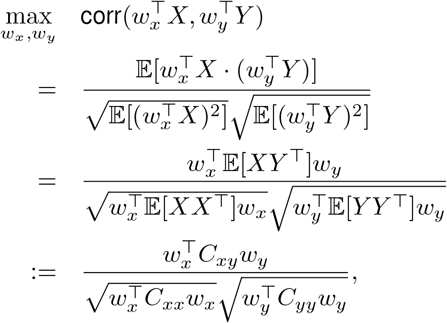

where *C*_*xx*_, *C*_*yy*_, and *C*_*xy*_ denote the covariance matrices of *X, Y*, and their cross-covariance, respectively. The expectation 𝔼 [·] is taken over the *n* samples.

We can fix the denominators by imposing the normalization constraints:

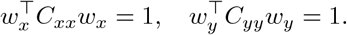

Additionally, we enforce the decorrelation constraint:

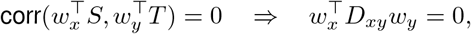

where *D*_*xy*_ := 𝔼 [*ST* ^*T*^] is the cross-covariance of *S* and *T*.

The final constrained optimization problem becomes:

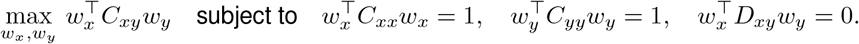

### Lagrangian Formulation

This is a non-convex quadratic problem, which is generally NP-hard. However, the structure of our problem allows for partial tractability. We introduce Lagrange multipliers *λ*_1_, *λ*_2_, and *λ*_3_ and define the Lagrangian:

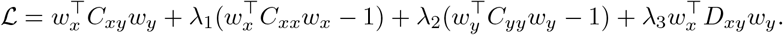

Taking derivatives and setting them to zero yields:

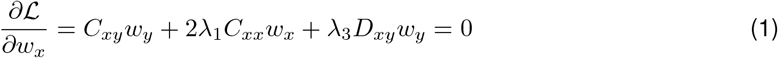

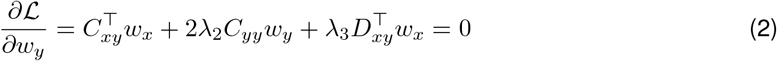

Multiplying Eq. (1) by 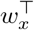 and Eq. (2) by 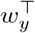, and subtracting, we find:

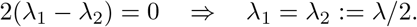

Substituting back, we obtain:

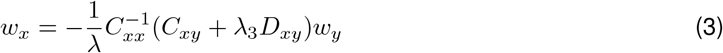

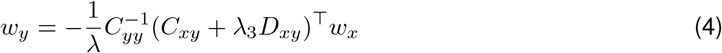

Combining these yields a generalized eigenvalue problem:

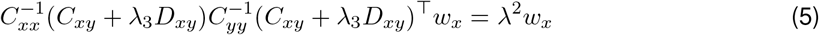

with the constraint:

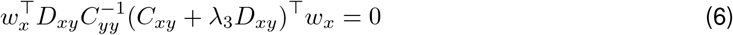

Both have to be solved at the same time for some *λ*_3_.

### Numerical Considerations

To solve Eqs. (5) and (6), we analyze limiting cases. Consider the regime where *λ*_3_ is large such that *C*_*xy*_ can be ignored relative to *λ*_3_*D*_*xy*_. Then:

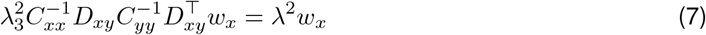

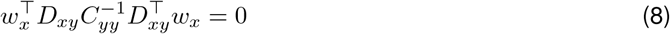

Multiplying the Eq. (7) by 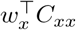 yields:

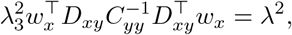

which contradicts Eq. (8) unless the left-hand side is zero. Thus, large *λ*_3_ do not yield valid solutions.

We define a critical threshold 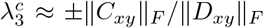, where ‖ · ‖_*F*_ denotes the Frobenius norm. For ease of computation, the Frobenius norm is a strategic choice. Valid solutions must lie within this range, and we can perform an exhaustive search over 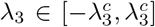 to find feasible solutions. In the following implementation, and the main text, we will drop the transposed symbol …^*T*^ for brevity.

### Numerical Solver

#### Algorithm 1

Full solver for the CRM optimization problem.

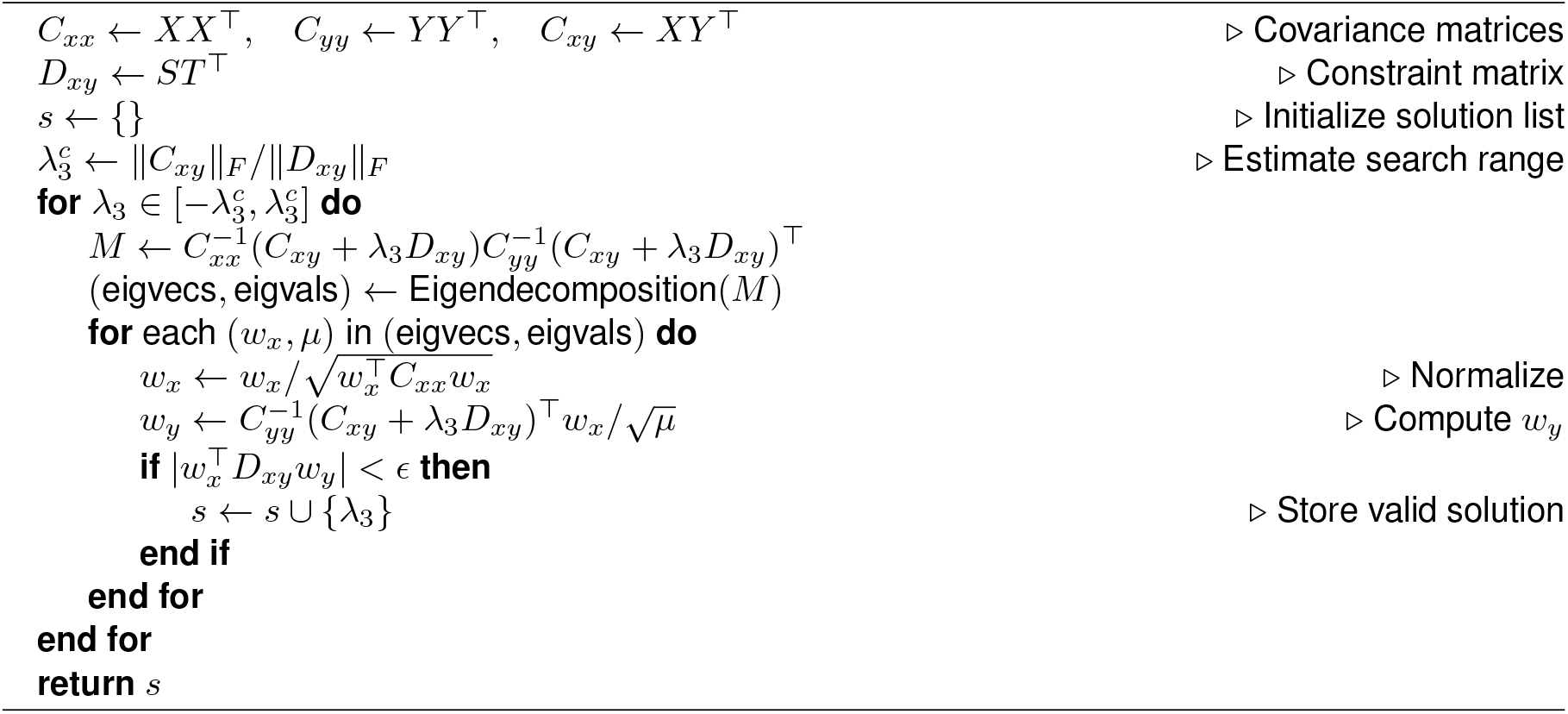

We provide an implementation of this scheme in Matlab as

~~~
 compute_weights_full.m
~~~

As this search is exhaustive, scanning through all values of *λ*_3_ can be computationally expensive. If only a single solution is desired, the problem can be reformulated as a root-finding task. Specifically, we seek the value of *λ*_3_ for which the constraint function 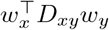 evaluates to zero. This leads to the following helper function, that can be solved with typical root finders.

#### Algorithm 2

Simplified solver for CRM using root-finding.

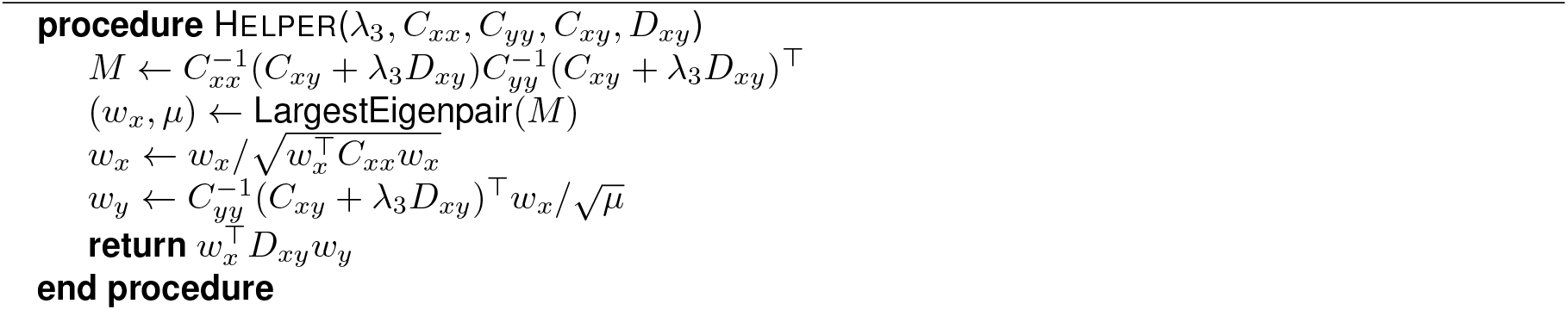

We provide an implementation of this simpler scheme in Matlab as

~~~
 compute_weights.m
~~~

### Regularization

In the limit where *D*_*xy*_ → 0, Eq. (5) reduces to:

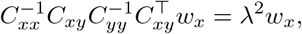

which corresponds to classical Canonical Correlation Analysis (CCA). Thus, CRM generalizes CCA by introducing a regularization term *λ*_3_*D*_*xy*_.

When *λ*_3_ → 0, the constraint in Eq. (6) is ignored, and the solution converges to standard CCA. Conversely, when *C*_*xy*_ *≈* 0, the Eigendecomposition becomes unstable. In such cases, the regularization term *λ*_3_*D*_*xy*_ can improve numerical stability.

In addition, it may be desirable to add regularization to the covariance matrices *C*_*xx*_ and *C*_*yy*_ which can further prevent overfitting^30^. I.e., we derive the regularized covariance matrices 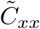 and 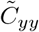, with

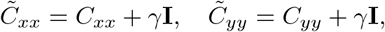

where *γ* is a small constant and **I** is the identity Matrix. We then apply algorithm 2 to the regularized covariance matrices.

### Principle of CRM on simulated data

This section describes Fig. 2. The entire figure was produced programmatically, and can be reproduced by running:

~~~
 Fig2_simulation.m
~~~

### CRM on numerous observations of variates

For panels a, b and c, we produced a list of *X* ∈ ℝ^*p×n*^ from normally distributed random numbers with unit variance, and computed from them *Y* = *M × X* +*α η* where *M* ∈ ℝ^*q×p*^ is a fixed matrix with random entries drawn from a normal distribution with unit variance. We chose *p* = *q* = 10 and *n* = 10000. The contribution of normally distributed noise *η* ∈ ℝ^*q×n*^ was set with a numerical parameter *α*. This procedure yielded the correlated datasets *X* and *Y*. In addition to these two datasets, we simulated another two random datasets *S* ∈ ℝ^*q×n*^*∼*𝒩 (*µ* = 0, *σ* = 1) and *T* = (*βN* + (1 − *β*)*M*) *× S* where *N* ∈ ℝ^*q×p*^*∼*𝒩 (*µ* = 0, *σ* = 1) is a random matrix, and *M* is taken from above. The partial alignment of *X, Y* and *S, T* via the shared matrix *M* makes the CRM problem challenging. The parameter *β* tunes this alignment, and was set ad-hoc to *β* = 1*/*4. For each noise level between *α* = 0.05 and *α* = 10 in steps of *Δα* = 0.5. *N* = 10 repeats were computed. Shown in Fig. 2a,b are means and SEM over these simulations. For the CCA comparison, the constraint was set to *D*_*xy*_ → 0. For panel c, we show the result for a single dataset from noise level *α* = 10, in which we scanned through *λ*_3_ in steps of *Δλ*_3_ = 0.01, and compute the 10 eigenvalues numerically using MATLAB’s eig() function.

### Removing correlated noise in the time domain

For panels d, e and f, we simulated a typical neurophysiological experiment. When recording neural activity with electrodes in two brain areas, signals can be contaminated with shared noise, for example line noise. This is captured by this simple model for data in brain areas *X* and *Y* :

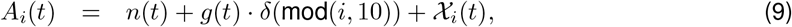

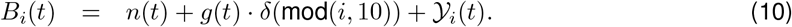

Here, *i* ∈ (1, …, *N*) is the electrode index for *N* = 100 electrodes in our simulation. *g*(*t*) is the signal or representation of interest. Enforced by the Kronecker delta function, it is only present if mod(*i*, 10) == 0. In other words, the signal of interest is only present on channels 0, 10, 20, …, 100. In addition, *n*(*t*) is a corrupting noise signal active on all electrodes such as line noise. Finally, both data contain uncorrelated noise content that is unique to each electrode and time. These are assumed to be stationary over time, and denoted as 𝒳_*i*_(*t*) and 𝒴_*i*_(*t*). In practice, we model 𝒳, 𝒴 to be normally distributed white noise, the signal of interest *g*(*t*) is a low frequency square wave function, and the noise signal *n*(*t*) = *A* sin(2*πf*_*n*_*t*) is a high frequency sinusoid. This produces the datasets *A, B* from with the covariances are computed as above. To use CRM, we also compute band-pass filtered versions of these data,

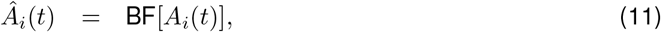

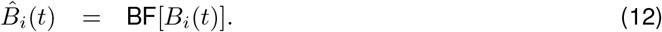

Here, BF() indicated a linear band-pass filter around the noise frequency *f*_*n*_. For simplicity we used MATLAB’s default band-pass filter, which is a minimum order-filter with a stop-band attenuation of 60dB and a steepness of 0.85. These filtered signals are used as *S, T* to compute the covariance *D*_*xy*_. Running CCA on these data produces uniform weights across electrodes. CRM on the other hand identifies the shared signals hidden on channels 0, 10, 20, … and ignores the other channels.

### Context-free alignment of neural representations and vector embeddings

#### Data set

We analyzed data from the “Podcast dataset”^25^ consisting of nine ECoG participants implanted with a total of 1, 330 electrodes. Patients listened to “Monkey in the Middle”, a 30-min podcast that appeared on “This American Life” (Chavis, 2017). The dataset further entails a transcript of the podcast, time-stamps for every word in the podcast, and the corresponding word-embeddings derived from various Large Language Models (LLMs), notably the Whisper model^24^, a model trained on 680, 000 hours of acoustic recordings to transcribe natural conversations.

#### Neural data processing

All analyses were realized on the pre-processed high frequency power data that are provided with the “Podcast” data set. The data were down-sampled to 100 Hz and epoched to word-onsets with intervals starting 4s before and ending 4s after word onset. Neural correlates of words were analyzed starting at word 20 from the beginning of the podcast.

#### Embedding processing

Word embeddings for each word in the story were reduced to 50 principal components^23^. For alignment, we matched each word-embedding to the corresponding word-epoch from the intracranial data. For the control time-series, we matched the word-embedding of the previous word to each word-epoch from the intracranial data.

#### Cross-validated alignment

The neural and behavioral data were partitioned for a 10-fold cross-validation. In each of 10 iterations, 90 percent of the data were used for computation of the alignment transformation and 10 percent of the data were held out to assess the correlation between projected components. Projection of the held-out data was done with the weights obtained from the training data. Alignment was computed across words (startig from word 20) and it was repeated for each time-point around word-onset. I.e, for the iEEG data at lag *X*_*l*_ ∈ ℝ^*c×n*^ (where *c* denotes the number of channels and *n* denotes the number of words), and the 50-d embedding data *Y*_*t*_ ∈ ℝ^50*×n*^ we obtained the covariance matrices *C*_*xx*_ ∈ ℝ^*c×c*^, *C*_*yy*_ ∈ ℝ^50*×*50^, and *C*_*xy*_ ∈ ℝ^*c×*50^. To compute CRM, the control covariance matrix *D*_*xy*_ ∈ ℝ^*c×*50^ was obtained from the covariance between the iEEG data *X*_*l*_ ∈ ℝ^*c×n*^, and the time-shifted embedding data *Y*_*t*−1_ ∈ ℝ^50*×n*^. To compute CCA, we simply defined *D*_*xy*_ := 0. For CCA and CRM we obtained projection vectors *w*_*x*_ and *w*_*y*_ via Algorithm 2 and computed the correlation in the held-out data as 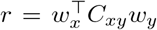. Correlations were then averaged across all test sets; grand-averages and standard error of the mean (SEM) were computed for CRM and CCA across patients.

#### Component loadings

For each fold and alignment method, we computed component loadings to quantify the contribution of individual iEEG channels to the extracted component at lag zero. We defined the component time series on the test data as 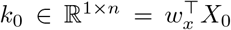 and computed the loading for channel (*i*, ∈ {1, …, *c*}) as the Pearson correlation between the channel’s time series at lag zero *x*_*i*,0_ ∈ ℝ^1*×n*^ and the component time-series. I.e., the loading for channel (*i*) was computed as the Pearson correlation between *x*_*i*,0_ and *k*_0_.

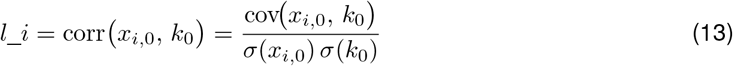

These channel–component correlations *l*_*i*_ were averaged across folds to yield a topographical map of loadings for CCA and CRM.

### Factorizing overlapping representations into their constituent parts

#### Data set

We re-analyzed a subset of data from Reagh et al^21^: In the video-viewing of their experiment, 20 subjects repeatedly watched 8 different videos fo 35s durations in 3 different runs. The 8 videos display 2 characters (Tommy and Lisa) in 2 different schematic situations (cafe and grocery stores) that are visited in 2 different contexts each (cafe 1 and 2, grocery store 1 and 2).

#### Data analysis

We analyzed the beta-weights that were obtained by the authors from a general linear model (GLM) fit on all runs using the AFNI 3dDeconvolve function. We focused on a region of interest in the medial Prefrontal Cortext (mPFC), where overlapping neural representations had been reported in the original study. In a first analysis, we correlated the vectors of beta weights in the mPFC between all 24 trials for each subject to obtain the Representational Similarity Matrix (RSM) of the raw data. We then computed the voxel-by-voxel cross-covariance matrix between all pairs of trials that formed part of the same schema (cafe, grocery), *L*_*xy*_ ∈ ℝ^*v×v*^ context (cafe 1, cafe 2, grocery store 1 and grocery store 2), *M*_*xy*_ ∈ ℝ^*v×v*^, and situation (e.g., Tommy in cafe 1 or Lisa in grocery store 2), *N*_*xy*_ ∈ ℝ^*v×v*^. We further computed the covariance between voxels within videos *C*_*xx*_ ∈ ℝ^*v×v*^ = *C*_*yy*_ ∈ ℝ^*v×v*^ and regularized the matrix at *γ* = 0.001. We then factorized representations by applying algorithm 2 to obtain 20 components where the correlations between trials of interest is maximized while the correlation of a confounding variable is controlled. Specifically, we first obtained 20 context-components by maximizing the correlation between all pairs of trials that formed part of the same context 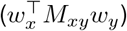, while keeping correlations between pairs of trials that formed part of the same schema (but not the same context, i.e., cafe 1 and cafe 2, and grocery store 1 and grocery store 2) at zero, 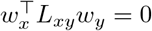.

Next, we obtained 20 situation-components by maximizing the correlation between all pairs of trials that formed part of the same situation 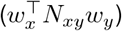, while keeping correlations between pairs of trials that formed part of the same context (but not the same situation, e.g., Tommy in cafe 1 and Lisa in cafe 1) at zero, 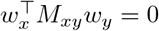.

Finally, we computed the correlations across the components between all trials for context and situation components.

#### Model comparison

To quantify the fit to various representational similarity models, we correlated each Representational Similarity Matrix (RSM) with a schematic, a contextual, and an episode model matrix. The model matrix contained one for every entry that was part of the model (i.e., same schema comparison, same context comparison, same episode comparison) and zero for every entry that was not part of the model (i.e., different schema, different context, different episode). Entries on the diagonal of the matrices (comparisons of trials to themselves) were excluded from this correlation. For the raw data (were correlations had not been fit to correspond to a representational structure of interest), we tested the model fit statistically with a dependent sample t-test across subjects; all p-values correspond to one-sided tests comparing the schema model to the other models and comparing the context model to the episode model.

### Unsupervised discovery of coupled spectral features in LFP recordings

This section describes Fig. 5. The entire figure was produced programmatically, and can be reproduced by running:

~~~
Fig5_analysis.m
~~~

We re-analyzed a previously published dataset^26^ from implanted wire arrays in the Hippocampus and medial prefrontal cortex in freely moving rats. The original recordings were made to study Fetal alcohol spectrum disorders in rats by administering alcohol during the first two postnatal weeks, and assessing the animals’ working memory through behavior in a maze. The analyzed dataset was taken from the control group. While a rat was solving the spatial working memory problem, recordings sampled at 2000 samples per second were made. The recordings were around 40 min long. We first z-scored the voltage traces, and then computed the spectrographic time-frequency representation *s*(*f, t*). We used a Hamming window to divide the signal into 1 second segments. The spectrum was then squared to compute power: |*s*(*f, t*) |^2^. To account for the typical 1/f decay, we computed the log-power for use with CRM: *X* = log(|*s*_HPC_(*f, t*) |^2^) and *Y* = log(|*s*_mPFC_(*f, t*) |^2^).

To select the noise band (i.e., band-pass filter the data), we applied a Gaussian-shaped filter along the frequency direction of the datasets. We chose *f*_0_ = 60 Hz and *s* = 10 Hz

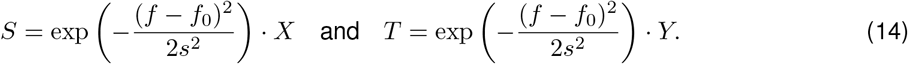

To aid numerical stability, the resulting *D*_*xy*_ was linearly rescaled to have roughly the same magnitude as *C*_*xy*_ using the Frobenius norm (see above). Finally, for easier visualization, the CRM and CCA components were mildly smoothed across the nearest 10 data points in space to produce panels Fig. 5e and Fig. 5f. No smoothing was applied to the other panels.

For the statistical test, we computed the Pearson correlation coefficient between the CRM components in Fig. 5e and Fig. 5f, and the distance of the animal from the home box *r* = corrcoef(CRM, *d*). We used the Euclidean distance 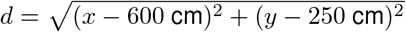. This number was compared to the null hypothesis that that there is no relationship between the two. The associated null distribution was computed with *N* = 10000 random shifts between the position and the CRM time series, and computing the correlation coefficient between the control data. The null distribution was unimodal, and centered around 0.

## Data availability

Data analyzed in this paper were provided by third parties. The iEEG data are publicly available in the podcast dataset^25^, fMRI and rodent data are available from the original authors upon request^21,26^.

## Code availability

All analysis code used in this manuscript is available from a github repository: https://github.com/s-michelmann/crm). The code version used for all figures is githash b6452a7.

## Acknowledgments

The authors wish to thank Zach Reagh and Amy Griffin for generously sharing their data, Ariel Goldstein for helpful discussions, and Ricardo Alejandro for helpful comments on an earlier version of this manuscript. MS is supported by a Burroughs Wellcome Fund’s Career Award at the Scientific Interface.

## Notes

### Competing Interest Statement

The authors have declared no competing interest.

